# Treatment of Hydrocephalus by Decreasing Inflammatory Cytokine Response Using GIT 27

**DOI:** 10.1101/2022.09.28.509923

**Authors:** Mira Zaranek, Carolyn Harris

## Abstract

**Background:** Surgical insertion of a ventricular shunt initiates a cytokine response shown to play a role in shunt failure caused by obstruction. These pro-inflammatory and anti-inflammatory cytokines cause astrocytes, amongst others, to enter an activated state which causes an increase in attachment. 4,5-Dihydro-3-phenyl-5-isoxazoleacetic acid (GIT 27) is a reagent with immunomodulatory properties which acts by blocking the main signaling protein on astrocytes and microglia called toll-like receptor 4 (TLR-4).

**Methods:** In this experiment, we tested the effect of GIT 27 on astrocytes when used as a pre-treatment, simultaneous treatment, and post-treatment relative to shunt insertion represented by the introduction of IL-1β or IL-10. Control, DMSO vehicle control, and GIT 27 treated sample groups were assayed for cell counts and cytokine concentration data.

**Results:** Exposure of astrocytes to suspended GIT 27 in a DMSO vehicle caused a decrease in cell attachment and a significant decrease in the concentration of the majority of cytokines. Comparisons of GIT 27 exposure times, represented by pre-, simultaneous, and post-treatment groups, showed that pre-treatment with GIT 27 is most effective at decreasing cellular attachment where post-treatment was generally the most effective at decreasing pro-inflammatory cytokine concentrations. In future practice, this could be embodied by pharmacologic dosing prior to shunting and/or slow release from the shunt surface.

**Conclusions:** GIT 27 is most effective at decreasing cell counts and cytokines when in-suspension compared to when attached to the shunt surface. Our data show that GIT 27 has the potential to be used as an effective way to modulate the cytokine response associated with shunt insertion.

## Introduction

Imbalance of cerebrospinal fluid (CSF) production and absorption often necessitates insertion of a ventricular shunt in hydrocephalus patients [1–4]. Treatment using a shunt system to divert excess CSF has extremely high failure rates, with 30% failing within one year post surgery and 85% within 10 years [5–8]. Obstruction of the proximal catheter holes with microglia, macrophages, and astrocytes make up 70% of these shunt failures [4,8–12]. While newly emerging evidence suggests that some shunt obstruction is dependent on periventricular tissue contact, our biorepository of obstructed ventricular catheters show us gross variability across samples, such as luminal obstruction, clotting, and evidence of proliferation, suggesting that both tissue contact and growth are factors [10,13].

Inflammatory response following shunt insertion plays a major role in the failure rates, specifically obstruction due to increased cell attachment [4]. Pro-inflammatory and anti-inflammatory cytokines are the main signaling chemicals of the immunoinflammatory response within the body [14]. Within the central nervous system, cytokines cause an increase in cell attachment due to the immunocompetent properties of astrocytes initiating astrogliosis [15]. Cytokines such as IL-1β, IL-6, IL-8, IL-10, MMP-7, and MMP-9 have all been shown to be upregulated in patients following a shunt insertion [14,15]. Central nervous system injury stemming from shunt insertion surgery transform astrocytes into its reactive phenotype [16]. Microglia and astrocytes contain a protein called toll-like receptor 4 (TLR-4), which aid in the regulation of the immune response by controlling signaling between cells [17,18].

Various methods have been developed to improve failure rates associated with shunt insertion by focusing on decreasing cell attachment and inflammation immediately surrounding the device responsible for the foreign body response. Utilizing coatings such as N-acetyl-L-cysteine, trypsin, and poly(2-hydroxyethyl methacrylate) have been shown to decrease cell adhesion [9,19,20]. These methods have shown the potential to decrease obstruction-dependent shunt failure. Other methods tested to regulate a more global inflammatory response utilize compounds such as polydopamine, hyaluronic acid, and decorin [21,22].

4,5-Dihydro-3-phenyl-5-isoxazoleacetic acid (GIT 27) is an immunomodulatory agent that works by reducing the production of pro-inflammatory cytokines [23,24]. GIT 27 has been shown to target TLR-4, and in doing so interferes with the secretion of tumor necrosis factor alpha (TNF-α) and consequently decreases concentrations of interleukin (IL)-1β, IL-10, and interferon gamma (IFN-γ) [23]. Previously, it has been shown that 50 mg/kg of GIT 27, also known as VGX-1027, decreases these three cytokines by inhibiting TLR-4 which in turn significantly reduced edema when administered as a four hour post-treatment [24]. Astrocytes, macrophages, microglia, and neurons are major sources of TLR-4 [25].

In this experiment, we tested the response of astrocytes treated with GIT 27 by analyzing total cell counts and cytokine concentration measurements using an enzyme-linked immunosorbent assay (ELISA). GIT 27 was administered as a pre-treatment, simultaneous treatment, or post-treatment, with respect to catheter insertion. This insertion was represented by exposing the samples to either IL-1β or IL-10 in the culture media. This mimics the upregulation of cytokines associated with the insertion of a catheter [14,15]. We hypothesized that GIT 27 treatment will cause a decrease in both cell count and cytokine concentrations to determine its utility as a surface coating, additive imbedded to the silicone shunt catheter or injection to reduce failure rate due to obstruction.

## Methods and Materials

### PDMS Catheter Creation

Catheter samples were made by creating a polydimethylsiloxane (PDMS, silicone) solution, Sylgard 184 (Dow Corning), at a ratio of 10:1 elastomer to curing agent. This solution was then homogenized and placed into a desiccator to remove bubbles. Once all the air bubbles were removed, 200 μL of the PDMS solution was pipetted into a 24 well plate to create flat disk samples for cell culturing. These samples were then placed in the desiccator again to ensure were no bubbles in the disks and left for 48 hours to cure on a flat substrate.

### Media Creation

A media stock solution was created by combining astrocyte media (ScienCell), supplement kit, 10 mL fetal bovine serum (FBS), 5 mL penicillin/streptomycin, and 5 mL astrocyte growth serum, (ScienCell), and an additional 10 mL FBS (ScienCell). IL-1β and IL-10 were diluted to a concentration of 50,000 ηg/mL and 100,000 ηg/mL, respectively, in 0.01 M phosphate buffer solution (PBS, w/v). GIT 27 was dissolved in a dimethyl sulfoxide (DMSO) vehicle (Sigma-Aldrich) to a concentration of 20 mg/mL (w/v).

Media solutions for the different groups were made directly before each media change. Cytokines IL-1β or IL-10 were added to the media stock solution for a final concentration of 14.67 ηg/mL. GIT 27 was added for a concentration of 1000 μg/mL in the media stock solution for our experimental group. Since GIT 27 is only soluble in DMSO, which has cytotoxic properties, a vehicle treatment group was performed to determine if it influenced cell counts or cytokine concentrations. Our vehicle group was exposed to 50 μL/mL DMSO within the media stock solution, which is proportional to the amount in the GIT 27 treated group. Concentrations were then made for evaluation of dose and time dependency (Figure 1).

**Figure 1:**
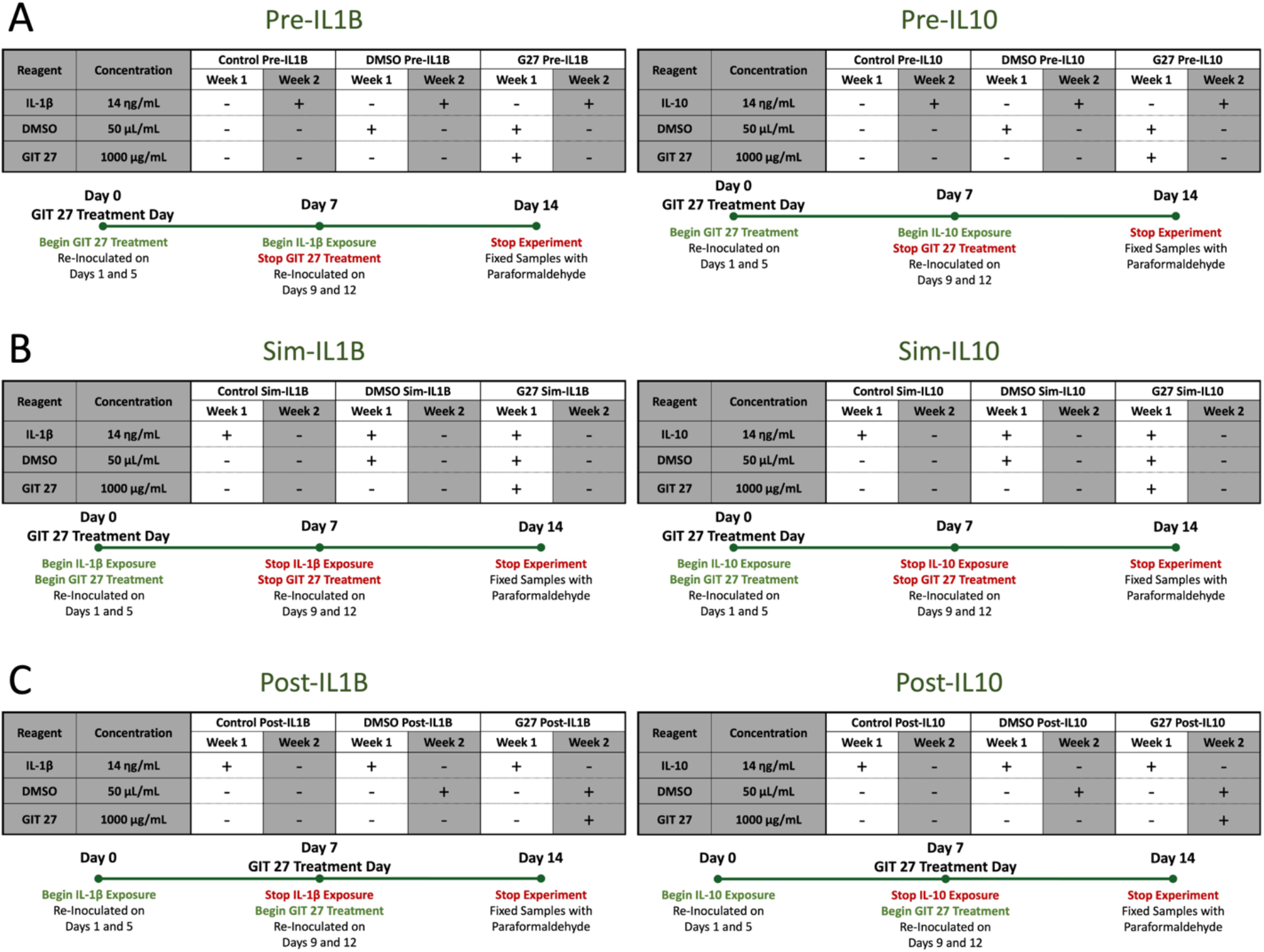
Visual representation of the treatment timelines and composition of media solution. A) Samples Exposed to Pre-Treatment B) Samples Exposed to Simultaneous Treatment C) Samples Exposed to Post-Treatment

### Cell Culture

Human astrocyte cells were initially grown in T75 flasks with media changes every three days using the media stock solution and washed with Hanks Balanced Salt Solution without Calcium and Magnesium (Gibco). Once the flasks came to confluency, the cells were trypsinized using Trypsin-EDTA at 0.25% (Gibco) and seeded at a seeding density of 250,000 cells per well. These samples were broken up into three treatment groups: control, DMSO, and GIT 27 tested as a pre-treatment, simultaneous treatment, and post-treatment (Figure 1). Each group and treatment time were then either exposed to IL-1β or IL-10 separately at concentrations described for media creation.

### GIT 27 Release Experiment

GIT 27 attachment to the surface of a shunt was completed using techniques similar to those described in previous work [26]. Briefly, ventricular shunts (Medtronic 27600) were cut into 1.5 cm segments and placed in the plasma etcher. Immediately after finishing the etching cycle, the samples were submerged in a solution made up of 2.5 mg/mL N-hydroxysuccinimide (NHS), 2.5 mg/mL 1-Ethyl-3-(3-dimethylaminopropyl) carbodiimide (EDC), and GIT 27 at either a 5, 50, and 1000 μg/mL concentration. Samples were left to incubate for 24 hours and then washed with 0.9% (w/v) saline solution. Samples were then incubated in 0.9% saline solution at 37°C to allow any GIT 27 to detach from the surface and erode into the saline solution.

At days 7, 14, 21, and 28, absorbance values were measured using the same methods as described in previous work [19]. Briefly, 2 μL of solution were placed on the nanodrop machine and measured using the a280 nanodrop setting on the Nano-Drop 2000 program. Absorbance values were recorded and compared to a standard curve made up of 7 concentrations evenly distributed between 0 and 1000 mg/mL of the GIT 27 in saline solution.

### Delivery Method

Following the nanodrop experiment, it was determined that the minimum concentration necessary to attach GIT 27 to the PDMS surface was 1000 μg/mL; at this concentration, attachment and release from the catheter was achievable. Analysis of the most effective delivery method was done by treating astrocytes with GIT 27 either attached to the surface PDMS or in suspension within the cell culture media. Using this concentration for sufficient surface coating, human astrocyte cells were cultured onto PDMS discs coated with GIT 27 or exposed to it in suspension within the media. Attached GIT 27 samples were conducted at concentrations 1000 and 3000 μg/mL, and for the in suspension concentrations 5, 50, and 1000 μg/mL in media stock solution were used.

Immunofluorescence was performed using glial fibrillary acidic protein (GFAP) and 4′,6-diamidino-2-phenylindole (DAPI) solution (ThermoFisher). GFAP primary and secondaries were diluted to a ratio of 1:1000 and 5:1000, respectively, in 0.4% Triton-X in PBS (w/v). Samples were incubated for 24 hours in each solution. After rinsing the samples with PBS, the samples were incubated in a 1:1000 DAPI solution in 0.4% Triton X solution for 30 minutes. Samples were then imaged submerged in PBS using a confocal microscope at 20× magnification and analyzed using Imaris software.

### Cell Count Collection

Images were taking using an inverted microscope (Fisher Scientific), 20× magnification, and microscope attachable digital camera following each media change on days 1, 5, 7, 9, 12, and 14. These images were taken at approximately the center of each 24 well plate well with an area of 5.9788 mm^2^. Counts were obtained using FIJI software by converting the images to 32-bit. Contrast was enhanced to the value of 0.5 and background was subtracted to a value of 10. Finally, the threshold was set to 23-70 to obtain the cell count of each image.

### ELISA

As previously published, multiplex assays according to the manufacturer’s protocol were performed by the Bursky Center for Human Immunology & Immunotherapy Programs Immunomonitoring Laboratory at Washington University School of Medicine [27]. Samples of culture media analyzed were collected on days 1, 5, 7, 9, 12, and 14 of culture following seeding day. Each frozen supernatant media sample was rapidly thawed at 37°C and centrifuged at 15,000 G for five minutes prior to incubating with the two-multiplex immunoassay for the following inflammatory cytokines: IL-1α, IL-1β, IL-6, TNF-α, IL-8, IL-10 (ThermoFisher Scientific), C3, and C1q (Millipore Sigma). Magnetic beads and assay buffer were added to all the wells of a 96 well-plate. Media samples and standards were then added in duplicate. The wells were thoroughly washed, and the detection antibody was then added followed by a streptavidin phycoerythrin incubation. Beads were resuspended with sheath fluid and 50 beads per region were acquired on a Luminex FLEXMAP3D system. The concentration of each analyte was then calculated by comparing the sample mean fluorescent intensity to the appropriate standard curve. Belysa v.1 software (Millipore Sigma) was used to generate a 5-parameter logistical curve fit algorithm. Protein concentration is reported as ρg/mL for each analyte.

### Statistical Analysis

Samples were exposed to three treatment groups, three treatment times, and exposure to either IL-1β or IL-10, with a total of 16 different samples tested with an n=5 replicates. The three treatment groups tested were a control, vehicle control (DMSO), and GIT 27 (denoted G27). Each treatment group was tested as a pre-treatment (denoted Pre-), dose simultaneous to treatment (denoted Sim-), and post-treatment (denoted Post-) as time points with respect to shunt insertion represented by exposure to either IL-1β (denoted IL1B) or IL-10 (denoted IL10). Utilizing the abbreviations described here we combined these to allow for an easier understanding of the results presented. An example of this can be control Pre-IL1B, DMSO Pre-IL1B, and G27 Pre-IL1B which represents the three treatment groups when used as a pre-treatment to IL-1β exposure.

Cell counts were analyzed for every sample, total of 80 replicates, from the images taken following each media change (Table 1). Cytokine concentrations were obtained for a total of n = 56 samples, Control 1 n = 4, Control 2 n = 4, DMSO n = 24, GIT 27 n = 24. Each sample was measured for IL-1β, IL-6, IL-8, IL-10, TNF-α, IL-1α, and IFN-γ concentrations. Concentrations for IFN-γ are not included since there was no detectable concentration in any of the treatment groups (Table 1).

**Table 1:**
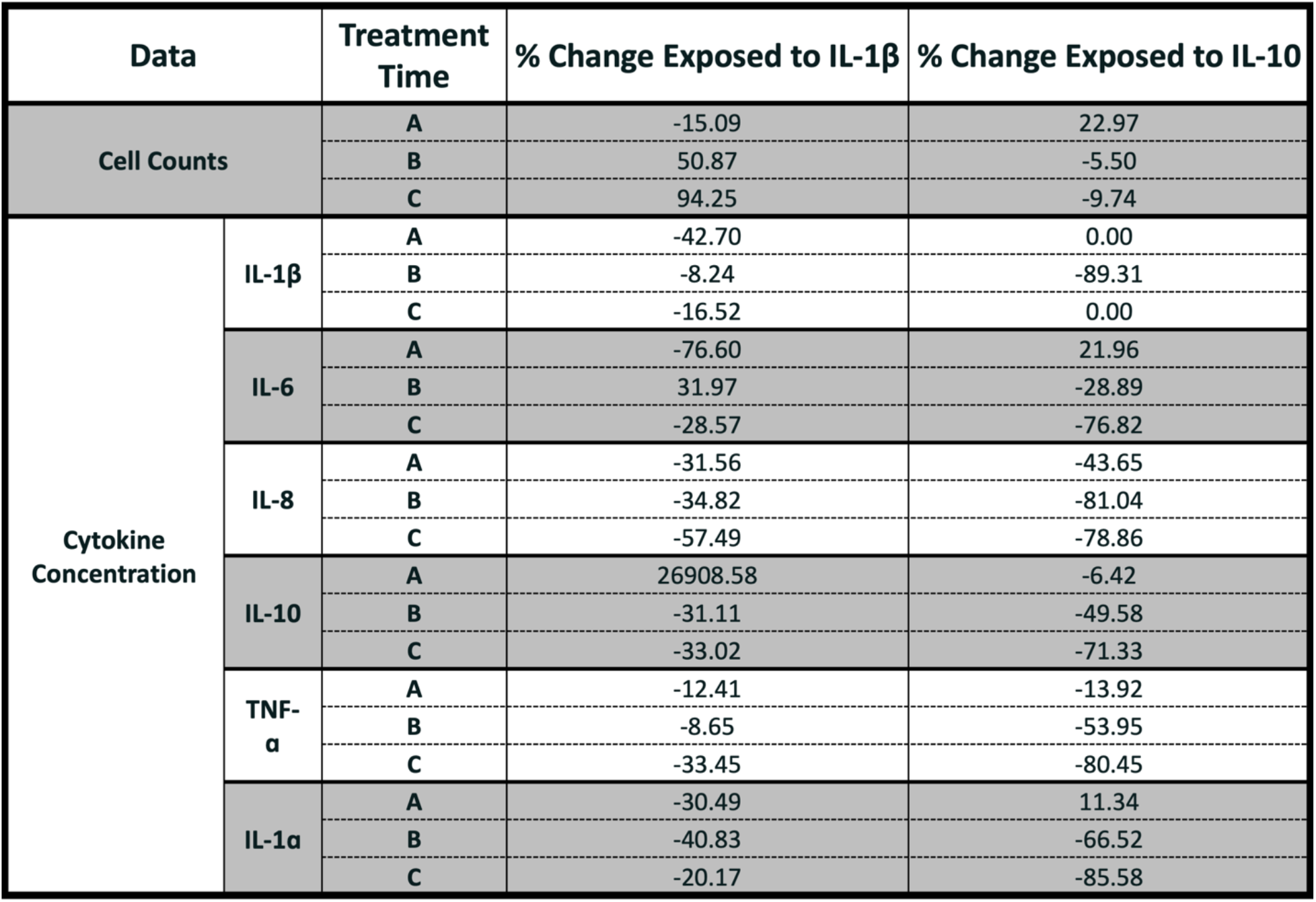
Percent change of average cell count and cytokine data for the GIT 27 treated samples compared to the DMSO treated vehicle. A) Samples Exposed to Pre-Treatment B) Samples Exposed to Simultaneous Treatment C) Samples Exposed to Post-Treatment

| Data | | Treatment Time | % Change Exposed to IL-1β | % Change Exposed to IL-10 |
| --- | --- | --- | --- | --- |
| Cell Counts |  | A | -15.09 | 22.97 |
|  |  | B | 50.87 | -5.50 |
|  |  | C | 94.25 | -9.74 |
| Cytokine Concentration | IL-1β | A | -42.70 | 0.00 |
|  |  | B | -8.24 | -89.31 |
|  |  | C | -16.52 | 0.00 |
|  | IL-6 | A | -76.60 | 21.96 |
|  |  | B | 31.97 | -28.89 |
|  |  | C | -28.57 | -76.82 |
|  | IL-8 | A | -31.56 | -43.65 |
|  |  | B | -34.82 | -81.04 |
|  |  | C | -57.49 | -78.86 |
|  | IL-10 | A | 26908.58 | -6.42 |
|  |  | B | -31.11 | -49.58 |
|  |  | C | -33.02 | -71.33 |
| | TNF-α | A | -12.41 | -13.92 |
|  |  | B | -8.65 | -53.95 |
|  |  | C | -33.45 | -80.45 |
| | IL-1α | A | -30.49 | 11.34 |
|  |  | B | -40.83 | -66.52 |
|  |  | C | -20.17 | -85.58 |

Normality was tested for both cell count and cytokine concentrations using the Anderson Darling equation for comparisons of treatment group and treatment time. Linear regression of the cell count data over time was used to determine time dependence. Nonparametric analysis of the treatment group and treatment time of the cell counts and cytokine concentrations using the Kruskal Wallis Test with the Dunn’s test with Bonferroni post-hoc test with α= 0.05. One-way ANOVAs followed by a Tukey post-hoc test with α = 0.05 was used for parametric results.

## Results

### Transient Nature of Cell Counts

Analysis using a linear regression of average cell count over the 14-day time period revealed a significant change in slope compared to the x-axis, slope of zero, for G27 Post-IL1B (*P* = 0.001) and DMSO Post-IL10 (*P* = 0.001) samples. These two groups reach a maximum average cell count on day 5 and can be observed to be fully confluent in the images. Observations of the images taken on day 7 showed either a peeling or release of cells from the surface, and when compared to day 5 the average cell count is lower. The release or peeling of cells in these samples occurred between days 5 and 7 when they were yet to be treated with any DMSO or GIT 27. All other samples expressed no significant change in slope.

### Comparison of Cell Counts by Treatment Group

The treatment groups exposed to IL-1β expressed a significant difference for all three treatment times: Pre-IL1B (*P* = 0.006), Sim-IL1B (*P* = 0.001), and Post-IL1B (*P* = < 0.0001) (Figure 2-4). G27 Pre-IL1B treatment group expressed a significantly lower cell count value when compared to control Pre-IL1B and insignificantly lower compared to DMSO Pre-IL1B groups (Figure 2). When comparing DMSO Sim-IL1B and DMSO Post-IL1B treatment groups there was a significantly lower number of cells compared to their associated control and G27 treated samples (Figure 3-4). Pre-IL10 samples showed a significant difference *(P* = 0.003) between treatment groups (Figure 2-4). DMSO Pre-IL10 had a significantly lower cell count when compared to control Pre-IL10 and insignificantly lower than G27 Pre-IL10 (Figure 2). There was no significant difference between samples under Sim-IL10 and Post-IL10 treatment conditions (Figure 3-4).

**Figure 2:**
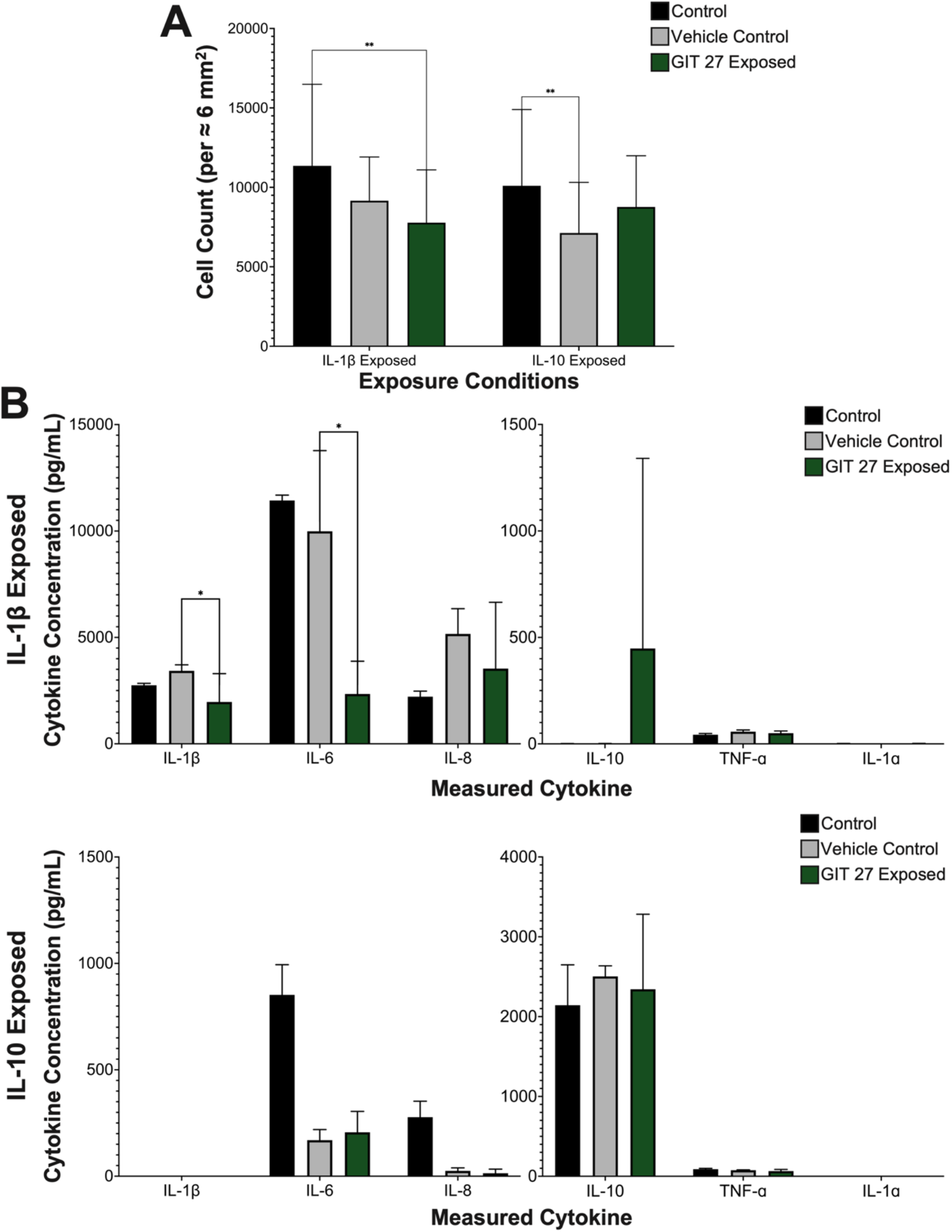
Comparisons of samples from the three treatment groups when under pre-treatment conditions. A) Cell Counts of Samples Exposed to a Pre-Treatment of GIT 27 B) Cytokine Concentrations of Samples Exposed to a Pre-Treatment of GIT 27

**Figure 3:**
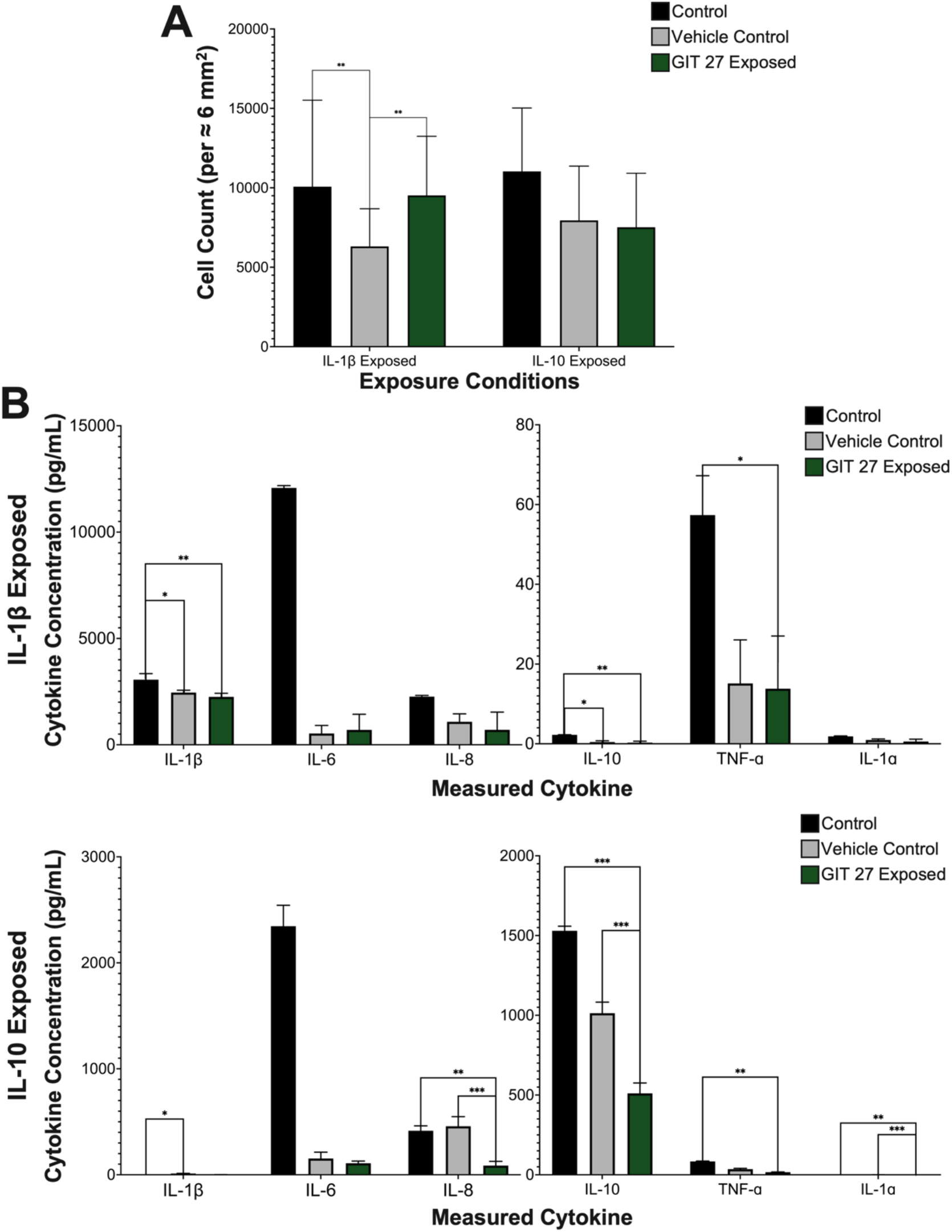
Comparisons of samples from the three treatment groups when under simultaneous treatment conditions. A) Cell Counts of Samples Exposed to a Simultaneous Treatment of GIT 27 B) Cytokine Concentrations of Samples Exposed to a Simultaneous Treatment of GIT 27

**Figure 4:**
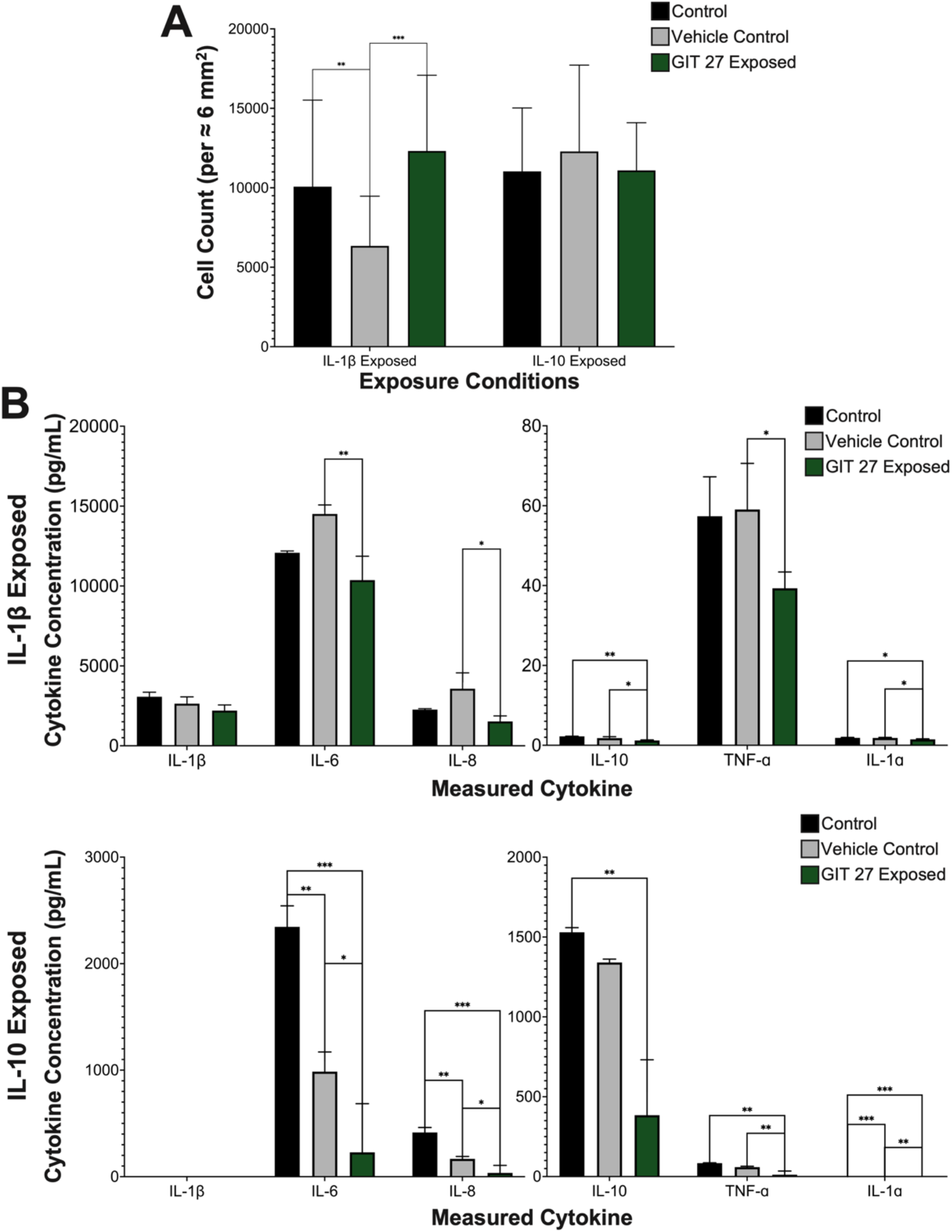
Comparisons of samples from the three treatment groups when under post-treatment conditions. A) Cell Counts of Samples Exposed to a Post-Treatment of GIT 27 B) Cytokine Concentrations of Samples Exposed to a Post-Treatment of GIT 27

### Comparison of Cytokine Concentrations by Treatment Group

Pre-IL1B treatment groups expressed a significant difference in IL-1β (*P* = 0.032) and IL-6 (*P* = 0.036) concentrations (Figure 2). The IL-1β and IL-6 concentration for G27 Pre-IL1B is significantly lower than DMSO Pre-IL1B, but not when compared to the control Pre-IL1B. Concentrations of IL-8, IL-10, TNF-α, and IL-1α expressed no significant difference for Pre-IL1B treatment groups. Samples treated under Pre-IL10 conditions expressed no significant difference for any of the cytokine concentrations (Figure 2).

Cytokine concentrations of Sim-IL1B treatment groups expressed a significant difference in concentration of IL-1β (*P* = 0.003), TNF-α (*P* = 0.008), and IL-1α (*P* = 0.026) (Figure 3). The concentration of IL-1β and IL-10 is significantly lower for DMSO Sim-IL1B and G27 Sim-IL1B compared to control Sim-IL1B. IL-1α concentration are significantly lower when treated with G27 Sim-IL1B compared to the control Sim-IL1B. Sim-IL10 treatment groups reported a significant difference in IL-1β (*P* = 0.019), IL-8 (p < 0.001), IL-10 (p < 0.001), TNF-α (*P* = 0.020), and IL-1α (p < 0.001) concentrations (Figure 3). IL-1β concentrations is significantly lower for the control Sim-IL10 compared to DMSO Sim-IL10, and IL-1α concentration is higher compared to G27 Sim-IL10. The concentration of IL-8 and IL-1α are significantly higher for DMSO Sim-IL10 and control Sim-IL10 compared to G27 Sim-IL10 individually. G27 Sim-IL10 concentration of IL-10 were significantly lower compared to both DMSO Sim-IL10 and control Sim-IL10.

Under Post-IL1B conditions, there was a significant difference in cytokine concentrations of IL-6 (*P* = 0.003), IL-8 (*P* = 0.011), IL-10 (*P* =0.004), TNF-α (*P* = 0.034), and IL-1α (*P* = 0.010) (Figure 4). There was a significantly lower concentration of IL-6, IL-8, and TNF-α for G27 Post-IL1B samples compared to DMSO Post-IL1B. Cytokine concentrations of IL-10 and IL-1α were significantly lower for G27 Post-IL1B compared to both control Post-IL1B and DMSO Post-IL1B. The concentration of IL-6 (p < 0.001), IL-8 (p < 0.001), IL-10 (*P* = 0.020), TNF-α (*P* = 0.002), and IL-1α (p < 0.001) were significantly different for Post-IL10 treatment groups (Figure 4). The concentration of IL-6, IL-8, and IL-1α for G27 Post-IL10 samples are significantly lower compared to the control Post-IL10 and DMSO Post-IL10. There is also a significantly lower concentration of these cytokines for DMSO Post-IL10 samples compared to the control Post-IL10. Concentrations of IL-10 and TNF-α are significantly lower for G27 Post-IL10 compared to the control Post-IL10. In addition to this, there is also a significantly lower concentration of TNF-α for G27 Post-IL10 compared to DMSO Post-IL10.

### Comparison of Cell Counts by Treatment Time

Comparisons of treatment time for the control group exposed to either IL1B or IL-10 expressed no significant difference and therefore is not included in this paper. The vehicle control treatment group showed a significant difference between treatment times when exposed to both IL1B (p < 0.001) and IL10 (p < 0.001) (Figure 5). DMSO Sim-IL1B and DMSO Post-IL1B samples expressed significantly lower cell counts compared to DMSO Sim-IL1B, and DMSO Post-IL10 is significantly higher than DMSO Pre-IL10 and DMSO Sim-IL10. The GIT 27 treatment group expressed a significant difference in the cell count when exposed to both IL1B and IL10. G27 Pre-IL1B and G27 Pre-IL10 has significantly fewer cells compared to G27 Post-IL1B and G27 Post-IL10 respectively. There is also a significantly lower cell count for G27 Sim-IL10 samples compared to G27 Post-IL10.

**Figure 5:**
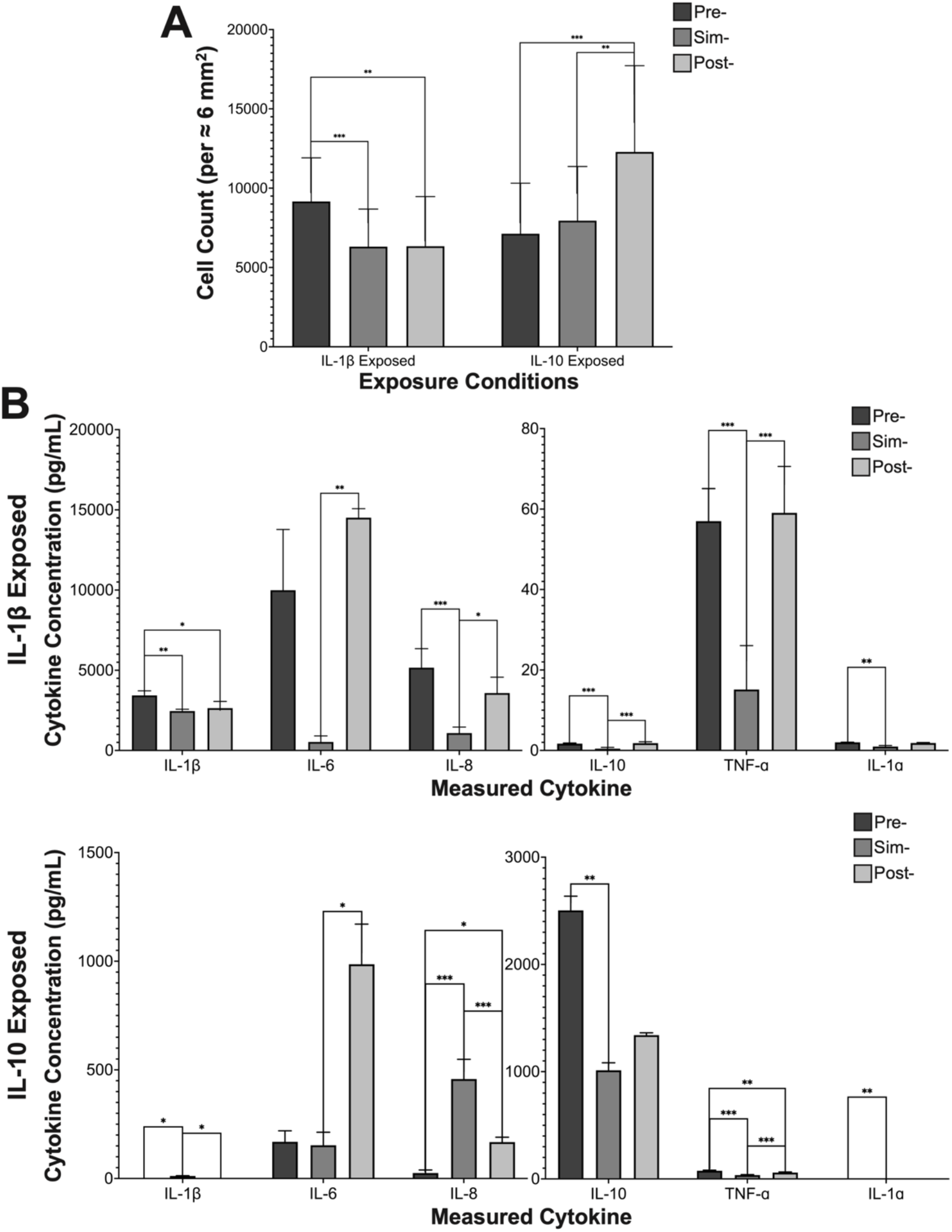
Comparisons of samples from the three treatment times when treated with the vehicle control DMSO. A) Cell Counts of Samples Exposed to DMSO at the Three Treatment Times B) Cytokine Concentrations of Samples Exposed to DMSO at the Three Treatment Times

### Comparison of Cytokines by Treatment Time

Treatment time comparisons of cytokine concentrations reported no significant difference for the control group exposed to IL1B or IL10 and is not included in this paper. Cytokine concentrations of the vehicle control group were significantly different for IL-1β (*P* = 0.003), IL-6 (*P* = 0.007), IL-8 (p <0.001), IL-10 (p <0.001), TNF-α (p <0.001), and IL-1α (p <0.001) when comparing the treatment times (Figure 6). DMSO Pre-IL1B has a significantly higher IL-1β concentration compared to both the DMSO Sim-IL1B and DMSO Post-IL1B. The concentration of IL-6 was significantly lower when under DMSO Sim-IL1B conditions compared to DMSO Post-IL1B, and DMSO Sim-IL1B is lower than the concentration of IL-1α for DMSO Pre-IL1B treatment times. DMSO Sim-IL1B has a significantly lower IL-8, IL10, and TNF-α concentration compared to both the DMSO Pre-IL1B and DMSO Post-IL1B treatment times.

**Figure 6:**
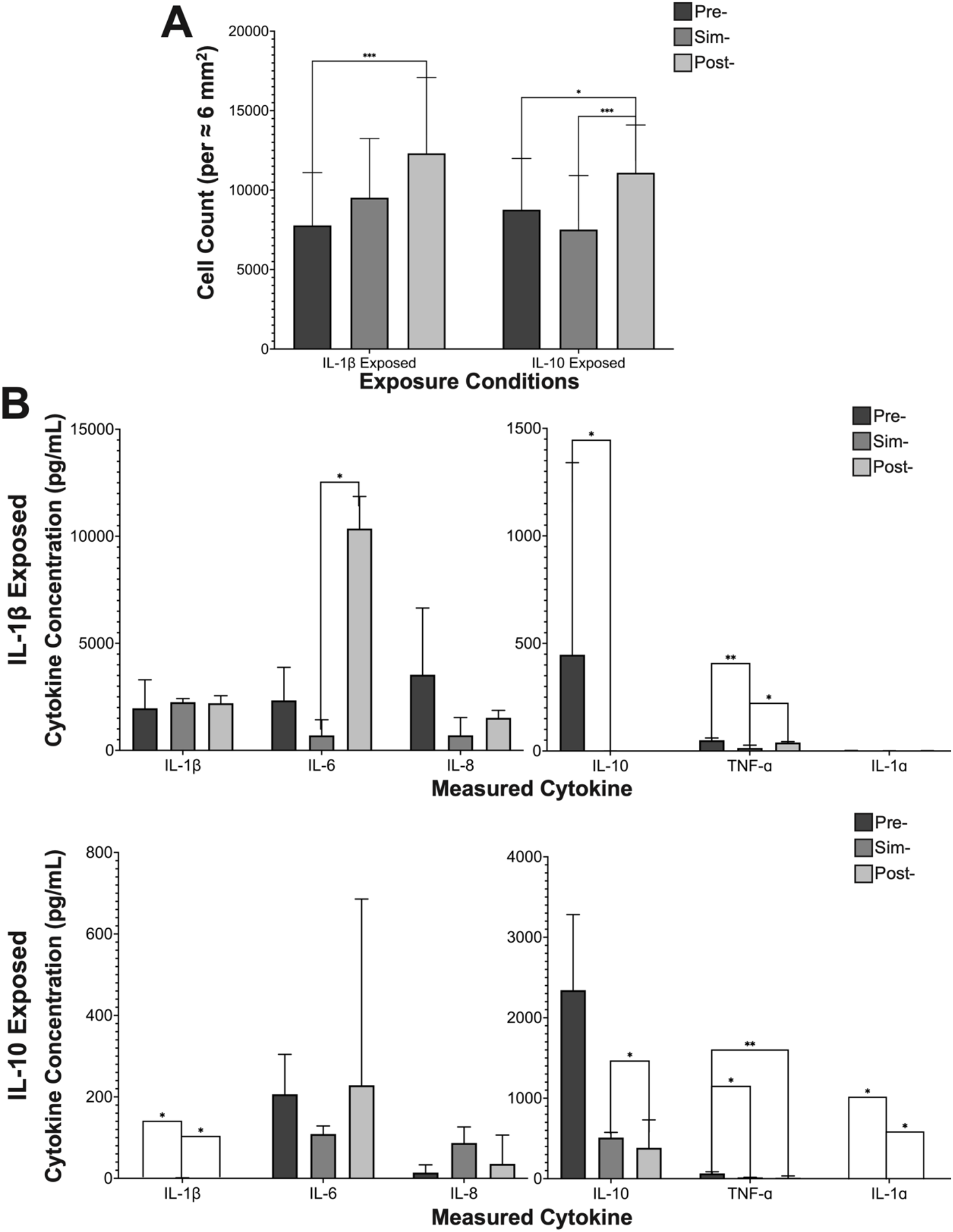
Comparisons of samples from the three treatment times when treated with the GIT 27. A) Cell Counts of Samples Exposed to GIT 27 at the Three Treatment Times B) Cytokine Concentrations of Samples Exposed to GIT 27 at the Three Treatment Times

GIT 27 treatment groups showed a significant difference in treatment time for cytokine concentrations of IL-6 (*P* = 0.015), IL-10 (*P* = 0.023), and TNF-α (*P* = 0.002) when exposed to IL1B, and when exposed to IL10 the concentrations IL-1β (*P* = 0.005), IL-10 (*P* = 0.018), TNF-α (*P* = 0.004), and IL-1α (*P* = 0.007) are significantly different (Figure 7). The concentration of IL-6 and TNF-α is significantly lower for G27 Sim-IL1B compared to G27 Post-IL1B, and lower for IL-10 and TNF-α compared to G27 Pre-IL1B. Concentrations of IL-1β and IL-1α are significantly higher for G27 Sim-IL10 compared to both G27 Pre-IL10 and G27 Post-IL10. G27 Post-IL10 concentration of IL-10 is significantly lower than the G27 Sim-IL10 group. TNF-α concentrations of G27 Pre-IL10 are significantly higher than the G27 Sim-IL10 and G27 Post-IL10 samples.

**Figure 7:**
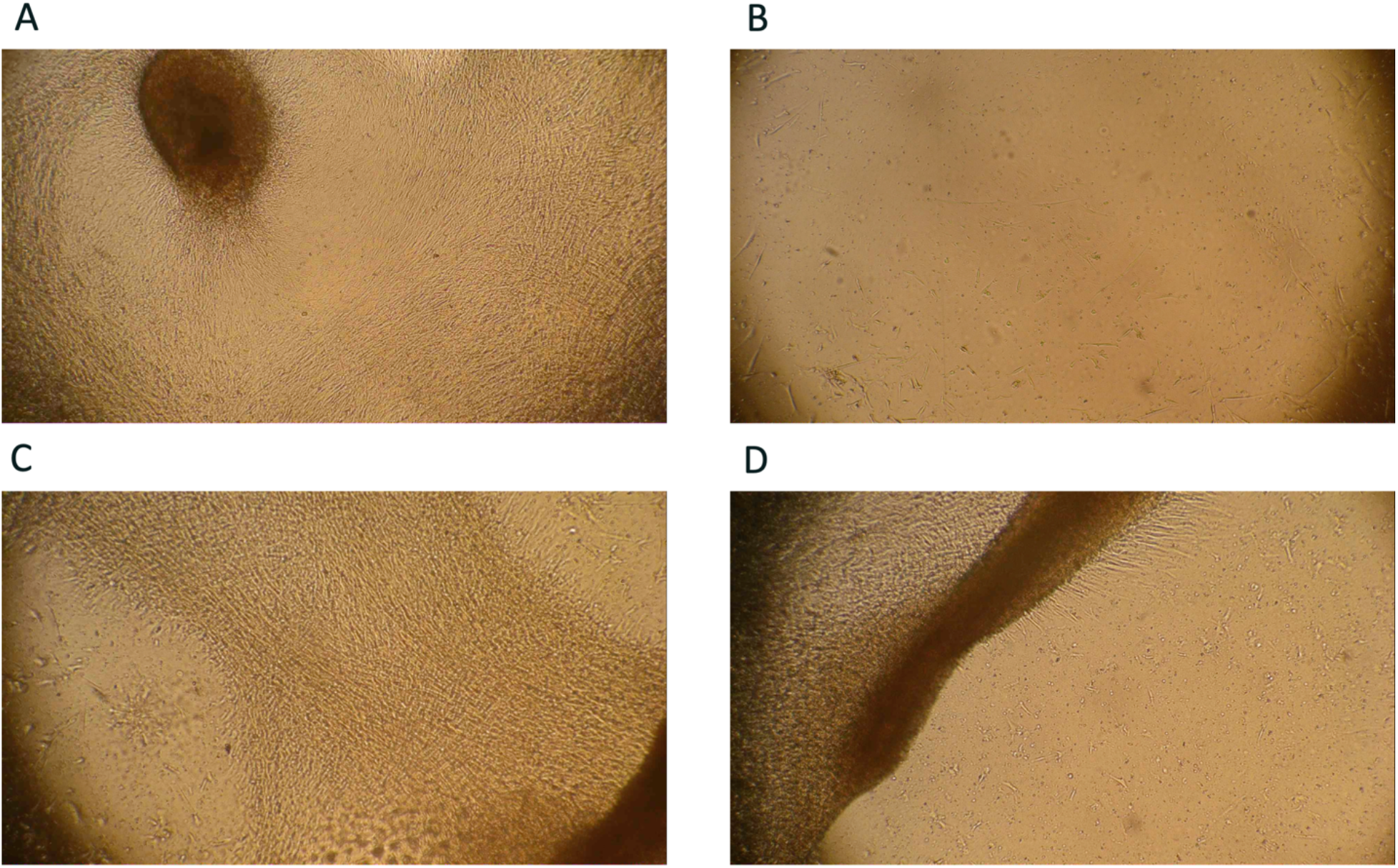
Visual example of samples that are confluent, bare, sluffing off, or peeling off. A) Confluent B) Bare C) Sluffing Off D) Peeling Off

## Discussion

To our knowledge, surface modification using GIT 27 is unexplored but has potential to serve as a pharmacologic supplement to treatment, as a released drug from the shunt catheter surface or as a surface chemical modification. As such, spectrophotometry was conducted to determine most effective delivery method/time and maximum concentration capable of coating a ventricular catheter. Determining this concentration, we ran an experiment to measure the absorbance values of the solution treated with 5 μg/mL, 50 μg/mL, and 1000 μg/mL of GIT 27. The lowest two concentrations, 5 μg/mL and 50 μg/mL, produced inconsistent values for both GIT 27 attached and released from the catheter. Data suggest that 1000 μg/mL GIT 27 following plasma etching physiosorbed the GIT 27 to the surface, of which was partially released over a four-week incubation period in saline.

Using this modification schema, we developed an in vitro model to determine the most effective GIT 27 delivery method, attached to PDMS surface or in suspension within the media, to decrease astrocyte attachment. Comparisons were made for the GIT 27 attached and in suspension samples to unmodified control, plasma etched control, vehicle control, and in suspension samples exposed to plasma etching. Qualitatively, analysis of the immunofluorescent total cell count (DAPI) and astrocyte cell bodies (GFAP) treated with a 1000 μg/mL concentration of GIT 27 in suspension exhibited the lowest amount of DAPI and GFAP labeling compared to all other groups. Samples exposed to plasma etching reported higher levels of GFAP compared to the samples that were not. Since GFAP attaches to cells within their cytoplasm this increase may indicate an increase of cell debris attached to the PDMS surface.

Initial experiments indicated that the concentration 1000 μg/mL of GIT 27 in suspension within cell culture media is the most effective treatment for decreasing astrocyte attachment. Utilizing this information, subsequent experiments focused on administration of GIT 27 in suspension versus bound to the catheter surface. The response of astrocytes when treated with GIT 27 at three time points was examined by analyzing cell count and cytokine concentration of astrocytes cultures. All treatment groups were exposed to either IL-1β or IL-10 to represent shunt insertion due to the significant increase in concentration during shunting [14,15]. We chose to expose our samples to IL-1β and IL-10 separately to generalize if the effect was more dependent on pro- or anti-inflammatory mediators, with an understanding of future work being necessary to distinguish cause and effect. Control samples were not treated with GIT 27 and only underwent exposure to the shunt insertion cytokines. Treatment time was investigated to understand if pre-treatment, simultaneous treatment, or post-treatment of GIT 27 in suspension, where cytokine exposure represents shunt insertion (Figure 1).

Results indicate that there is a significant time dependence for G27 Post-IL1B and DMSO Post-IL10, whereas all other groups have no time dependence. Qualitatively, we see these samples as overly confluent on day 5 then, on day 7, lacking cells after a large cell mass sloughed off the surface in these two groups (Figure 7). Since, for the majority of the samples, there is not time dependence, we were able to analyze the cell counts for the samples regardless of the time the counts were taken.

Cell count data for G27 Pre-IL1B reports a significantly lower number of cells compared to control Pre-IL1B. However, since the DMSO Pre-IL1B samples are insignificantly higher than G27 Pre-IL1B it cannot be determined that GIT 27 is the exclusive cause for the decrease (Figure 2a). Concentration IL-1β and IL-6 were significantly lowered for G27 Pre-IL1B compared to the DMSO Pre-IL1B (Figure 2b). Both of these cytokines are known to be associated with astrocyte activation, and these decreased levels could attribute to the insignificantly lower number of cells treated with GIT 27 [28]. Even though insignificant, there was a trend showing increased IL-1β in DMSO Pre-IL1B compared to the control Pre-IL1B. This was potentially caused by the cytotoxic properties of DMSO and was somewhat negated by GIT 27 exposure. Cell counts of DMSO Pre-IL10 are significantly lower than control Pre-IL10, but not G27 Pre-IL10, which indicates that any decrease in cell attachment is probably caused by the vehicle (Figure 2a). Under these conditions there was no significant change in the concentration of the measured cytokines.

Analysis of Sim-IL1B and Post-IL1B revealed that DMSO is the reason for any cell loss that had taken place under these conditions (Figure 3a and 4a). Concentrations of IL-1β and IL-10 are also significantly lower for DMSO Sim-IL1B and G27 Sim-IL1B when compared to control Sim-IL1B. The lower number of cells when treated with DMSO, for both Sim-IL1B and Post-IL1B, compared to the controls and GIT 27 treated could attribute to the reduced level of these cytokines for these samples. Since the number of cells from G27 Sim-IL1B and G27 Post-IL1B is not significantly different to their associated controls, it seems as though GIT 27 may have blocked the cytotoxic effect. TNF-α concentrations are significantly lower for G27 Sim-IL1B samples in relation to the control Sim-IL1B. Since GIT 27 interference of TLR-4 signaling pathways, it will subsequently cause a decrease in the production of TNF-α [23]. This decrease in TNF-α can also potentially contribute to the decrease in IL-1β concentrations for G27 Sim-IL1B as well.

DMSO Post-IL1B samples had an insignificant increase in cytokine concentration for IL-6, IL8, and TNF-α but G27 Post-IL1B significantly lowered this value. Again, this may indicate that DMSO exposure can further exacerbate the cytokine response, although minor, but when paired with GIT 27 it neutralizes this negative effect. G27 Post-IL1B caused the concentration of IL-10 and IL-1α to be significantly lower compared to both control Post-IL1B and DMSO Post-IL1B. Decreased levels of TNF-α indicates that GIT 27 is effectively blocking TLR-4 which subsequently decreased the other cytokine levels.

Sim-IL10 and Post-IL10 samples expressed no significant difference between cells counts for the three treatment groups, indicating GIT 27 has no effect on the cell count (Figure 3a and 4a). GIT 27 caused the concentrations of IL-8, IL-10, and IL-1α to all significantly decrease for G27 Sim-IL10. These samples also significantly lowered the concentration of TNF-α compared to the control. Blocking of TLR-4 with GIT 27 may cause a decrease in TNF-α production, and a decrease in IL-8 and IL-10 will cause less signaling between cells causing a lower cytokine response. G27 Post-IL10 caused levels of IL-6, IL-8, IL10, TNF-α and IL-1α to decrease. GIT 27 simultaneous treatment and post-treatment seems to have no effect on the cell count when exposed to IL-10 but can decrease the cytokine response.

Optimal treatment time of GIT 27 is an important aspect for the potential of using it to decrease the cytokine response. DMSO Pre-IL1B had the least amount of effect on cell count when compared to the other treatment times, indicating that the cytotoxic effect of DMSO is low when it’s a part of a pre-treatment. Under G27 Pre-IL1B conditions the cell count was significantly lower than Post-IL1B and insignificantly less than Sim-IL1B. Therefore GIT 27 is most effective when Pre-IL1B, and in combination with DMSO’s low effect as a pre-treatment it might be the best treatment time. DMSO Post-IL10 and G27 Post-IL10 caused the least amount of cell loss compared to the other treatment times. Since the other treatment time points had significantly lower cell counts for both DMSO and GIT 27 treated groups we cannot determine the best treatment time when exposed to IL-10.

Greatest impact of cytokine concentration occurred when under DMSO Sim-IL1B conditions. Lower numbers of cells under these conditions could potentially have led to the lower concentrations of cytokines. Comparisons between GIT 27 treated samples indicate a generally lower concentration of cytokines when Sim-IL1B, excluding IL-1β. DMSO Post-IL10 samples have increased levels of IL-6 and TNF-α which play a role in cell proliferation which probably can account for the significantly higher number of cells in Post-IL10. Further work will need to be done to determine the best treatment time for GIT 27 exposure.

## Conclusion

GIT 27 has immunomodulatory properties that interfere with the signaling process associated with TLR-4. In this study, we determined that the most effective delivery method for treatment using GIT 27 is in suspension, which demonstrated minimal cell attachment. In clinical practice, this may look like oral delivery or localized release from the shunt catheter surface. Analysis of the treatment groups showed some decrease in cell counts, but the mechanism for action is yet to be understood. DMSO seems to have the potential to play a factor in the lower numbers, even if this is minimal. Conversely, we have determined that GIT 27 causes a majority of the decrease in the overall cytokine response. Transiently, exposure of GIT 27 prior to shunting and/or an influx of IL-1β and IL-10 appears to be the most effective at reducing cell attachment on silicone shunt material. GIT 27 post-treatment seems to have the largest effect on cytokine concentration. Future work should be done exposing both microglia and astrocytes to GIT 27 and the PDMS material to create a more physiological cytokine environment.

## Acknowledgements

Research reported in this publication was supported by the National Institute of Neurological Disorders and Stroke of the National Institutes of Health under award number R01NS094570. Approximately 90% was financed with federal dollars. The content is solely the responsibility of the authors and does not necessarily represent the official view of the Hydrocephalus Association or the National Institutes of Health.

## Author Contributions

**Mira Zaranek:** Conceptualization, methodology, formal analysis, investigation, validation, data curation, data analysis, project administration, writing—original draft, visualization, and writing—review and editing.

**Carolyn A. Harris:** Conceptualization, methodology, resources, supervision, project administration, funding acquisition, and writing—review and editing.

## Data Availability Statement

Raw data were generated at Wayne State University and can be made available to other qualified research scientists for appropriate applications and collaborative efforts in accordance with Wayne State University & Close Curly Quote’s materials transfer guidelines as well as those set by NIH and the Hydrocephalus Association. Patentable work will be reviewed by the Wayne State Technology Transfer Office prior to data sharing, were applicable.

## References

1. Pople IK. Hydrocephalus and shunts: What the neurologist should know. Neurology in Practice. 2002;73.

2. Corns R, Martin A. Hydrocephalus. Surgery. 2012;30:142–8.

3. Leinonen V, Vanninen R, Rauramaa T. Cerebrospinal fluid circulation and hydrocephalus. 1st ed. Handb Clin Neurol. Elsevier B.V.; 2018.

4. Harris C, Pearson K, Hadley K, Zhu S, Browd S, Hanak BW, et al. Fabrication of three-dimensional hydrogel scaffolds for modeling shunt failure by tissue obstruction in hydrocephalus. Fluids Barriers CNS. BioMed Central; 2015;12:1–15.

5. Riva-Cambrin J, Kestle JRW, Holubkov R, Butler J, Kulkarni A V., Drake J, et al. Risk factors for shunt malfunction in pediatric hydrocephalus: A multicenter prospective cohort study. J Neurosurg Pediatr. 2016;17:382–90.

6. Shannon CN, Acakpo-Satchivi L, Kirby RS, Franklin FA, Wellons JC. Ventriculoperitoneal shunt failure: An institutional review of 2-year survival rates. Child’s Nervous System. 2012;28:2093–9.

7. Stone JJ, Walker CT, Jacobson M, Phillips V, Silberstein HJ. Revision rate of pediatric ventriculoperitoneal shunts after 15 years: Clinical article. J Neurosurg Pediatr. 2013;11:15–9.

8. Harris CA, Resau JH, Hudson EA, West RA, Moon C, Black AD, et al. Effects of surface wettability, flow, and protein concentration on macrophage and astrocyte adhesion in an in vitro model of central nervous system catheter obstruction. J Biomed Mater Res A. 2011;97 A:433–40.

9. Hanak BW, Hsieh CY, Donaldson W, Browd SR, Lau KKS, Shain W. Reduced cell attachment to poly(2-hydroxyethyl methacrylate)-coated ventricular catheters in vitro. J Biomed Mater Res B Appl Biomater. 2018;106:1268–79.

10. Hariharan P, Sondheimer J, Petroj A, Gluski J, Jea A, Whitehead WE, et al. A multicenter retrospective study of heterogeneous tissue aggregates obstructing ventricular catheters explanted from patients with hydrocephalus. Fluids Barriers CNS. BioMed Central; 2021;18:1–12.

11. Chittiboina P, Pasieka H, Sonig A, Bollam P, Notarianni C, Willis BK, et al. Posthemorrhagic hydrocephalus and shunts: What are the predictors of multiple revision surgeries? Clinical article. J Neurosurg Pediatr. 2013;11:37–42.

12. Gluski J, Zajciw P, Hariharan P, Morgan A, Morales DM, Jea A, et al. Characterization of a multicenter pediatric-hydrocephalus shunt biobank. Fluids Barriers CNS. BioMed Central; 2020;17:1–10.

13. Kraemer MR, Koueik J, Rebsamen S, Hsu DA, Salamat MS, Luo S, et al. Overdrainage-related ependymal bands: A postulated cause of proximal shunt obstruction. J Neurosurg Pediatr. 2018;22:567–77.

14. Harris CA, Morales DM, Arshad R, McAllister JP, Limbrick DD. Cerebrospinal fluid biomarkers of neuroinflammation in children with hydrocephalus and shunt malfunction. Fluids Barriers CNS [Internet]. BioMed Central; 2021;18:1–14. Available from: https://doi.org/10.1186/s12987-021-00237-4

15. Hyvärinen T, Hagman S, Ristola M, Sukki L, Veijula K, Kreutzer J, et al. Co-stimulation with IL-1β and TNF-α induces an inflammatory reactive astrocyte phenotype with neurosupportive characteristics in a human pluripotent stem cell model system. Sci Rep. 2019;9:1–15.

16. Deren KE, Packer M, Forsyth J, Milash B, Abdullah OM, Hsu EW, et al. Reactive astrocytosis, microgliosis and inflammation in rats with neonatal hydrocephalus. Exp Neurol. Elsevier Inc.; 2010;226:110–9.

17. Zacharowski K, Zacharowski PA, Koch A, Baban A, Tran N, Berkels R, et al. Toll-like receptor 4 plays a crucial role in the immune-adrenal response to systemic inflammatory response syndrome. 2006;

18. Pascual-Lucas M, Fernandez-Lizarbe S, Montesinos J, Guerri C. LPS or ethanol triggers clathrin-and rafts/caveolae-dependent endocytosis of TLR4 in cortical astrocytes. J Neurochem. Blackwell Publishing Ltd; 2014;129:448–62.

19. Al-Saloum S, Zaranek M, Horbatiuk J, Gopalakrishnan P, Dumitrescu A, McAllister JP, et al. Analysis of N-acetyl cysteine modified polydimethylsiloxane shunt for improved treatment of hydrocephalus. J Biomed Mater Res B Appl Biomater. 2021;109:1177–87.

20. Achyuta AKH, Stephens KD, Lewis HGP, Murthy SK. Mitigation of reactive human cell adhesion on poly(dimethylsiloxane) by immobilized trypsin. Langmuir. 2010;26:4160–7.

21. Xue P, Li Q, Li Y, Sun L, Zhang L, Xu Z, et al. Surface modification of poly(dimethylsiloxane) with polydopamine and hyaluronic acid to enhance hemocompatibility for potential applications in medical implants or devices. ACS Appl Mater Interfaces. 2017;9:33632–44.

22. Botfield H, Gonzalez AM, Abdullah O, Skjolding AD, Berry M, Mcallister JP, et al. Decorin prevents the development of juvenile communicating hydrocephalus. Brain. 2013;136:2842–58.

23. Stojanovic I, Cuzzocrea S, Mangano K, Mazzon E, Miljkovic D, Wang M, et al. In vitro, ex vivo and in vivo immunopharmacological activities of the isoxazoline compound VGX-1027: Modulation of cytokine synthesis and prevention of both organ-specific and systemic autoimmune diseases in murine models. Clinical Immunology. 2007;123:311–23.

24. Laird MD, Shields JS, Sukumari-Ramesh S, Kimbler DE, Fessler RD, Shakir B, et al. High mobility group box protein-1 promotes cerebral edema after traumatic brain injury via activation of toll-like receptor 4. Glia. 2014;62:26–38.

25. Shi H, Hua X, Kong D, Stein D, Hua F. Role of Toll-like receptor mediated signaling in traumatic brain injury. Neuropharmacology. Elsevier Ltd; 2019;145:259–67.

26. Harris CA, Resau JH, Hudson EA, West RA, Moon C, Black AD, et al. Reduction of protein adsorption and macrophage and astrocyte adhesion on ventricular catheters by polyethylene glycol and N-acetyl-L-cysteine. J Biomed Mater Res A. 2011;98 A:425–33.

27. Fatemeh Khodadadei1*, Rooshan Arshad2, Diego M. Morales3, Jacob Gluski4, Neena I. Marupudi4, James P. McAllister II3, David D. Limbrick Jr3, Carolyn A. Harris1, 4 5. The Role of A1/A2 Reactive Astrocyte Expression in Hydrocephalus Shunt Failure.

28. Aschner M. Immune and inflammatory responses in the CNS: Modulation by astrocytes. Toxicol Lett. 1998;102–103:283–7.

